# Identification of an N-acetylneuraminic acid-presenting bacteria isolated from a healthy human microbiome

**DOI:** 10.1101/2020.08.24.265504

**Authors:** Zhen Han, Peter S. Thuy-Boun, Wayne Pfeiffer, Vincent F. Vartabedian, Ali Torkamani, John R. Teijaro, Dennis W. Wolan

## Abstract

N-acetylneuraminic acid is the most abundant sialic acid in humans and is generally expressed as the terminal sugar on intestinal mucus glycans. Several pathogenic bacterial species harvest sialic acid from the mucus, diet, as well as other intestinal sources and display this sugar on their own surface to evade sialic acid-binding immunoglobulin-type lectin (Siglec)-mediated host immune surveillance. While previous studies have identified bacterial enzymes associated with sialic acid catabolism, no reported methods permit the selective labeling, tracking, and quantitation of sialic acid-presenting microbes within complex multi-microbial systems. Here, we apply an interdisciplinary approach combining metabolic labeling, click chemistry, metagenomic, and whole-genome sequencing to selectively track and identify sialic acid-presenting microbes from a cultured healthy human fecal microbiome. We isolated and identified a new strain of *Escherichia coli* that incorporates sialic acid onto its own surface. Analysis of the sequence data reveals that this *E. coli* strain encodes for the NanT, NeuA, and NeuS genes necessary for harvesting environmental sialic acid and generating the capsular polysialic acid. We envision that this method is applicable to the detection and quantitation of sialic acid-presenting bacteria from human, animal, and environmental microbiomes, as well as investigating the importance of other carbohydrates to commensal and pathogenic bacteria.

## Introduction

Several methodologies and model systems have been critical to understanding gut microbiome taxonomic composition and the influence these compositions exert on host physiology and intestinal disease pathology (1–3). Metagenomics and metabolomics as well as germ-free and gnotobiotic mouse models have helped establish how specific microbes and small-molecule metabolites modulate host responses to a variety of diseases (4–7). These methods, in combination with the genetic manipulation of specific bacterial strains, has revealed key catabolizing enzymes and pathogen-associated molecular patterns (PAMPs) associated with microbiome-related metabolism, host immune activation, and bacterial infection (8–10). Despite these methods and models, the ability to selectively label and track bacteria with specific catabolic functionalities in a highly complex and metabolically active microbiome remains limited.

N-acetylneuraminic acid (Neu5Ac, termed sialic acid/SA here) is one of over thirty known sialic acid analogs. It is the predominant form of sialic acid in humans and is presented as the terminal residue on surface-exposed glycans, glycoproteins, and glycolipids (11). SA is recognized by immunoinhibitory sialic acid-binding immunoglobulin-like lectins (Siglecs) to prevent autoimmunity (12). Interestingly, select human pathogenic microbes have evolved to express this human SA epitope on their own cellular surface to evade host immune surveillance and clearance (8,13–15). For example, group B streptococcus (GBS), a common cause of sepsis in human newborns, presents terminal α2,3-linked SA on its capsular polysaccharide (CPS) to bind Siglecs expressed by neutrophils, macrophages, and platelets and block immune activation (8,16,17). Sequencing analysis revealed that GBS uses a tripartite transporter to translocate environmental SA onto its CPS to augment this immune evasion (8,18). We posited that certain gut commensal organisms likely apply similar SA-mediated protective mechanisms to gain survival advantages. As such, the selective labeling and identification of these microbes from complex microbiome samples may reveal new SA-regulated host-microbe crosstalk mechanisms that could potentially be exploited for therapeutic development aimed at gut microbiome-related diseases.

Here, we report the application of an SA-based azide-containing probe N-acetyl-9-azido-9-deoxy-neuraminic acid (Neu5Ac9N_3_/Sia9N_3_), combined with flow cytometry and metagenomic and whole-genome sequencing, to selectively label, isolate, and identify SA-presenting bacteria from a complex cultured human fecal microbiome sample. Broadly, applications of metabolic probe labeling followed by click chemistry-tagging for visualization have been used to image and track pre-labeled bacterial components, including peptidoglycans and lipopolysaccharides (19–21). However, there is limited use of SA-based probes to label bacterial glycans, especially in a highly complex microbial sample. With respect to eukaryotic labeling, “clickable” SA-based probes and its metabolic precursor (*i.e.,* sialic acid-alkyne, sialic acid-azide, and mannosamine-alkyne) have been widely used for the study of sialoglycans in both *in vitro* and *in vivo* systems (22–26). We sought to expand the application of SA-based metabolic probes to selectively label SA-presenting bacteria from a distal gut microbiome. Fecal microbiome samples collected from a healthy human volunteer were cultivated in the presence of the Sia9N_3_ metabolic probe (27) and bacterial incorporation of Sia9N_3_ was examined by flow cytometry analysis following copper-free azide coupling with azadibenzocyclooctyne-conjugated biotin (ADIBO-BTN/DBCO-BTN) and streptavidin-Alexa 647 staining (28). We identified a healthy human fecal microbiome sample containing bacteria that can readily incorporate Sia9N_3_. With fluorescence activated cell sorting (FACS) and 16S rDNA sequencing analyses, we found that the Sia9N_3_-incorporating bacteria belonged to the *Escherichia* genus. Isolation of the SA-incorporating bacteria via colony screening with whole-genome sequencing analysis suggested that the isolated bacteria is a new strain of *E. coli* that likely employs the NanT-NeuA-NeuS system to integrate environmental sialic acid to form polysialic acid on its capsular polysaccharide structure (29,30).

## Results

### FACS-based screening helps identify microbiome constituents capable of Sia9N_3_ incorporation

We surveyed fecal samples collected from one healthy human volunteer over the course of three months. Fecal microbes were cultured in a Gifu media (31) supplemented with Sia9N_3_ at a physiologically relevant concentration (200 μM) (32) under anaerobic atmosphere for 20 h. Microbes were centrifuged, extensively washed, subjected to DBCO-BTN conjugation with streptavidin-Alexa 647, and examined for fluorophore incorporation with flow cytometry analysis (**Fig. 1**). Among three microbiota samples tested (designated H1, H2, and H3), more than 60% of the bacterial cells from one of the cultured human fecal samples (*e.g.,* H2) readily displayed fluorescence in response to Sia9N_3_ treatment (**Fig. 1**). Furthermore, a dose-dependent assay revealed that bacterial fluorescence labeling is dependent on concentrations of both Sia9N_3_ and DBCO-BTN. Use of 200 μM Sia9N_3_ in culture media with 50 μM DBCO-BTN for azide conjugation resulted in the maximum percentage of labeled bacterial cells (**Figs. 2A** and **B**). Further increases in the concentration of either labeling reagent only amplified the fluorescence intensity on the labeled bacterial cells without increasing background fluorescence or numbers of labeled cells, which suggested that Sia9N_3_ selectively labeled a subset of SA-catabolizing bacteria (**Fig. S1**).

**Fig. 1.**
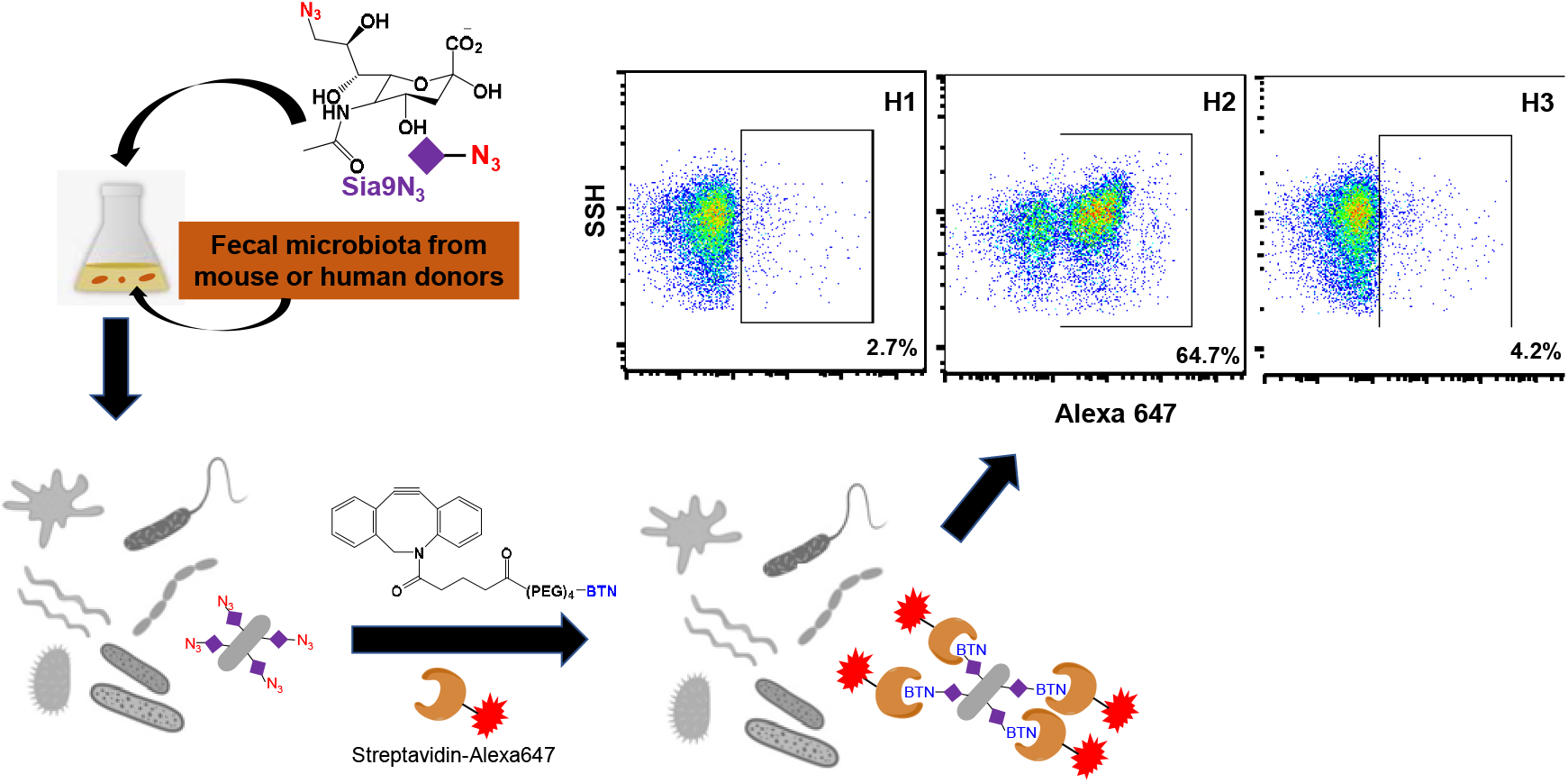
Metabolic labeling of SA-presenting microbes from a fecal microbiome sample. An interdisciplinary approach combining metabolic labeling, click chemistry, and flow cytometry revealed a human fecal microbiome sample that contains SA-presenting bacteria.

**Fig. 2.**
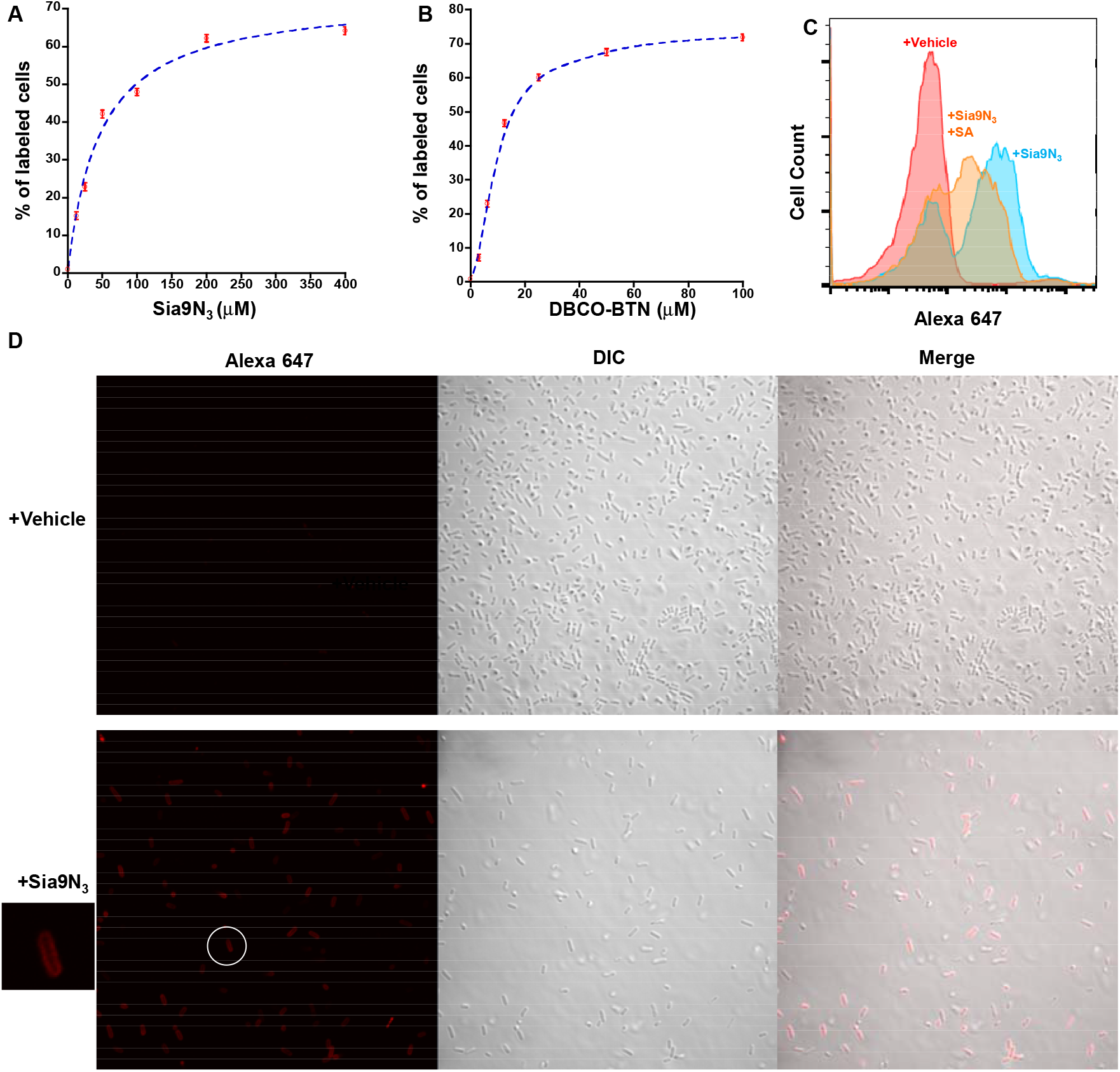
Characterization of Sia9N_3_ induced fluorescence staining with flow cytometry and confocal fluorescence microscopy. Labeling of SA-presenting microbes is Sia9N_3_ (**A**) and DBCO-BTN (**B**) dose-dependent. **C.** Addition of excess natural SA partially competed for fluorescence labeling. **D**. Confocal fluorescence microscopic imaging showed that Sia9N_3_ induced fluorescence labeling on the bacterial cell surface.

We next assessed if Neu5Ac would compete with Sia9N_3_ for surface expression by bacteria when supplemented into the 20 h cultures. Addition of excess Neu5Ac resulted in reduced labeling of bacteria by Sia9N_3_ (**Figs. 2C** and **S2A**), which suggested that bacteria may use the same or partially overlapped transporters and enzymes for SA and Sia9N_3_ transportation and catabolism. Additionally, these results indicate that Sia9N_3_ is biologically relevant as a metabolic probe since the substitution of hydroxyl to azide on C9 did not significantly affect its interaction with the SA-binding proteins. Labeled bacteria were also subjected to fluorescence microscopic analysis to visualize where the bacteria are labeled (*i.e.,* intracellular, periplasmic, cell wall, etc.). These experiments provided visual evidence that Sia9N_3_ or its azido-metabolites are prominently delivered to the bacterial cell surface (**Fig. 2D**). This finding is consistent with the observation that certain bacteria can present environmental SA onto its surface capsular polysaccharides and/or lipopolysaccharides (33).

### Bacterial surface Sia9N_3_ is removed by microbial sialidases

Many gut bacteria can express sialidases to release SA from sialoglycans, providing themselves or other bacteria with a source of carbon, nitrogen, and building blocks for cell wall biosynthesis (34,35). To determine if the Sia9N_3_ probe is presented on bacteria to form sialoglycans, we treated the metabolically labeled H2 cultured sample with a purified recombinant sialidase (BT0455, NCBI gene ID: 1071627) (36) from *Bacteroides thetaiotaomicron,* followed by sequential DBCO-BTN conjugation, streptavidin-Alexa 647 staining, and flow cytometry. Sialidase treatment dramatically reduced the number of fluorescently labeled bacterial cells and fluorescence intensity (**Figs. 3A** and **S2B**). This result suggested that the majority of bacterial fluorescent labeling is due to Sia9N_3_ transported into certain bacterial cells and presented onto the cell surface to form sialoglycans. The inability to completely eliminate fluorescence labeling is likely due to substrate specificity of BT-0455 that includes α2,3-, α2,6-, and α2,8-linked sialic acid substrates (36). Sia9N_3_ may be installed on glycans unrecognizable by BT-0455 and/or Sia9N_3_ is catabolized and surface expressed as a modified SA or other sugar.

**Fig. 3.**
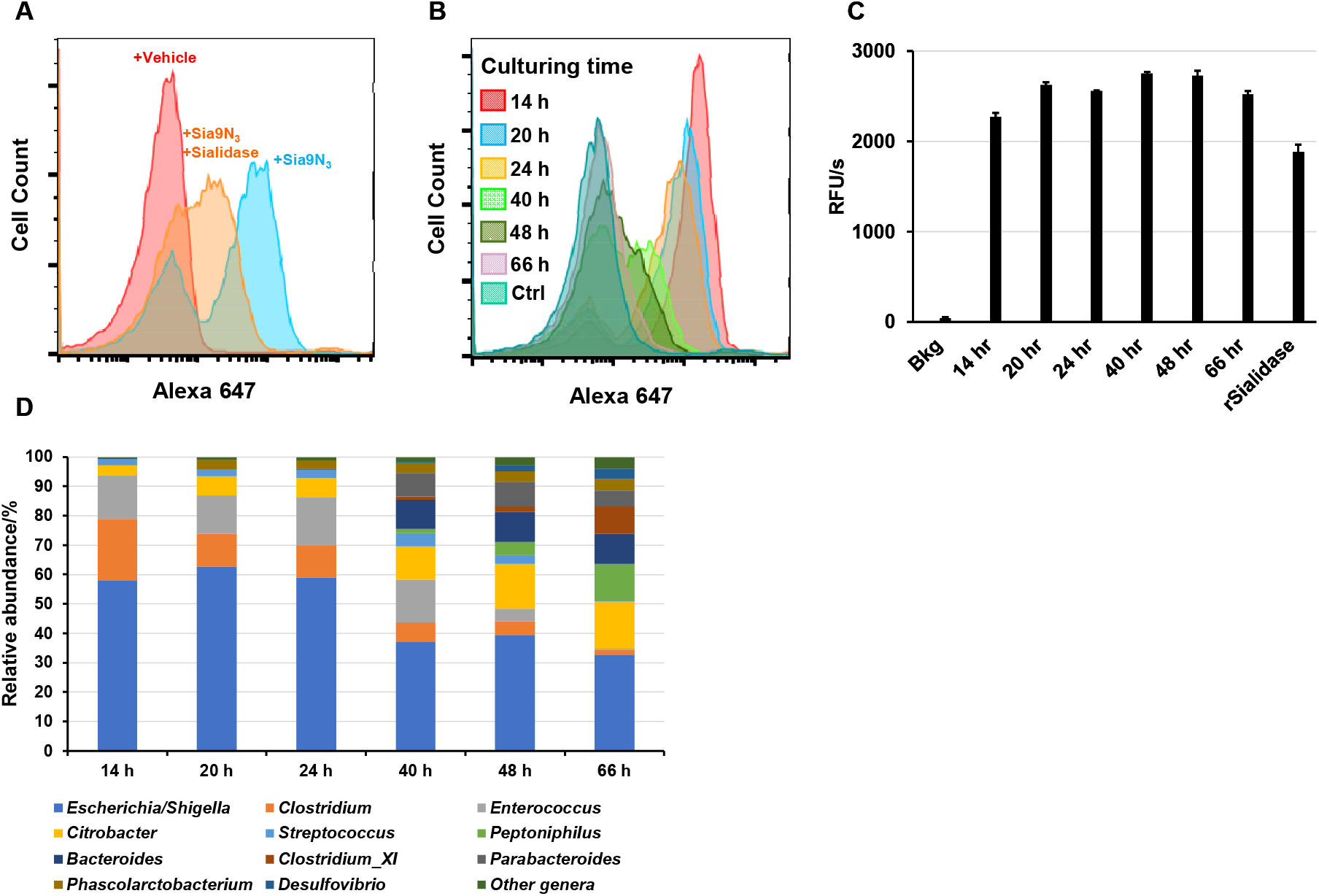
Bacterially incorporated surface Sia9N_3_ can be removed by microbial sialidases over time. **A.** Treatment of labeled microbes before DBCO-BTN conjugation with a recombinant sialidase significantly reduces fluorescence labeling. The incorporated Sia9N_3_ is removed by microbial sialidases over time as fluorescence labeling was gradually removed during microbiota cultivation (**B**) and microbial sialidase activity was constantly detected in the culture media, 500 pM recombinant BT0455 was used as a positive control (**C**). **D.** 16S rDNA sequencing revealed the change of taxonomic composition during H2 microbiome culture.

Sialic acid can be released from sialoglycans by microbial sialidases as demonstrated (**Fig. 3A**), which can promote the growth of certain SA-utilizing pathogens in the gut and lead to inflammatory diseases (32). To explore if Sia9N_3_ can also be released from the bacterially presented sialoglycans by other bacteria within the cultured sample, we tracked Sia9N_3_ incorporation over time during the plateau phase of bacterial growth. Briefly, the H2 fecal microbiome sample was grown with Sia9N_3_ for 14 to 66 h and the bacteria were quantitated for Sia9N_3_ incorporation by flow cytometry as described. Interestingly, detection of Sia9N_3_ labeling was maximally present at 14 h and gradually decreased until almost all labeling was eliminated by 40 h of incubation (**Fig. 3B**). Cultured media from each sample for every timepoint was assessed for sialidase activity, as measured by catalytic turnover of 4-methylumbelliferyl-N-acetylneuraminic acid (4-MUANA). Measurable sialidase activity was observed across all samples and suggests that bacterially secreted sialidases could remove Sia9N_3_ from bacterial sialoglycans over time (**Fig. 3C**). In parallel to the sialidase analysis, 16S rDNA sequencing showed that the compositional diversity of the cultured microbiota increased over time (**Fig. 3D**). At 14 h, the microbiome collection was dominated by *Escherichia/Shigella* (58% relative abundance), *Clostridium* (21%), and *Enterococcus* (15%). Common commensal organisms, including *Citrobacter, Peptoniphilus*, *Bacteroides*, and *Parabacteroides,* began to appear after 24 h of culture and their relative abundances peaked at 16%, 13%, 10%, and 5%, respectively, at 66 h of culture. The corresponding relative abundance of *Escherichia/Shigella*, *Clostridium*, and *Enterococcus* dropped to 33%, 2%, and 0%, respectively. Our results suggest that there is no correlation between Sia9N_3_ removal and abundance of secreted sialidase activity within the cultured media. While the bloom of other commensal bacteria at later time points may suppress the detection of Sia9N_3_-presenting microbes, we posit that the complete elimination of Sia9N_3_ detection is likely due to sialidase activity. The sialidases responsible for the removal of Sia9N_3_ may not be present at early time points and are introduced by late-blooming commensal organisms, such as *Bacteroides*, that appear at later time points, as determined by 16S rDNA analysis. Importantly, this shift in taxonomic composition may reflect protective mechanisms by which commensal bacteria target virulence factors expressed on pathogenic/pathobiont bacteria by exposing surface antigens for host recognition. Alternatively, commensal bacteria could be modifying environmental conditions to reduce the growth of pathogenic/pathobiont bacteria. All together, these data demonstrated that Sia9N_3_ incorporation by bacteria in culture is dynamically regulated by other bacteria, nutrients, and bacterial sialidases.

### The isolated Sia9N_3_-presenting bacteria is a new strain of *Escherichia coli*

We next separated the Sia9N_3_ incorporating bacteria from a 20-h culture of the H2 microbiome sample with FACS. The isolated fraction that yielded a high fluorescence signal due to Sia9N_3_ incorporation was subjected to 16S rDNA sequencing and revealed that the *Escherichia* genus was significantly enriched post sorting (**Figs. 4A** and **B**). We next subjected the cultured and primary H1-H3 microbiome samples to 16S rDNA sequencing to determine if differences in the bacterial taxonomic composition accounted for the selective labeling of the H2 microbiome sample only. Of note, 20 h of culture significantly altered the composition of all samples, and therefore, made the Sia9N_3_-presenting bacteria detectable in H2 as the *Escherichia* genus was significantly enriched compared to the starting fecal sample (**Fig. S3**). In addition, although only H2 contains Sia9N_3_-presenting bacteria, *Escherichia* dominated all three cultured microbiome samples as the most abundant genus. These results suggest that *Escherichia* presented in the H1-3 cultured microbiome samples may differ at the strain level as only the *Escherichia* in H2 has the catabolic genes to incorporate Sia9N_3_. An additional possibility is that the isolated *Escherichia* organism from H2 is present across all samples; however, the SA-presenting genes are not activated in the cultured H1 and H3 samples.

**Fig. 4.**
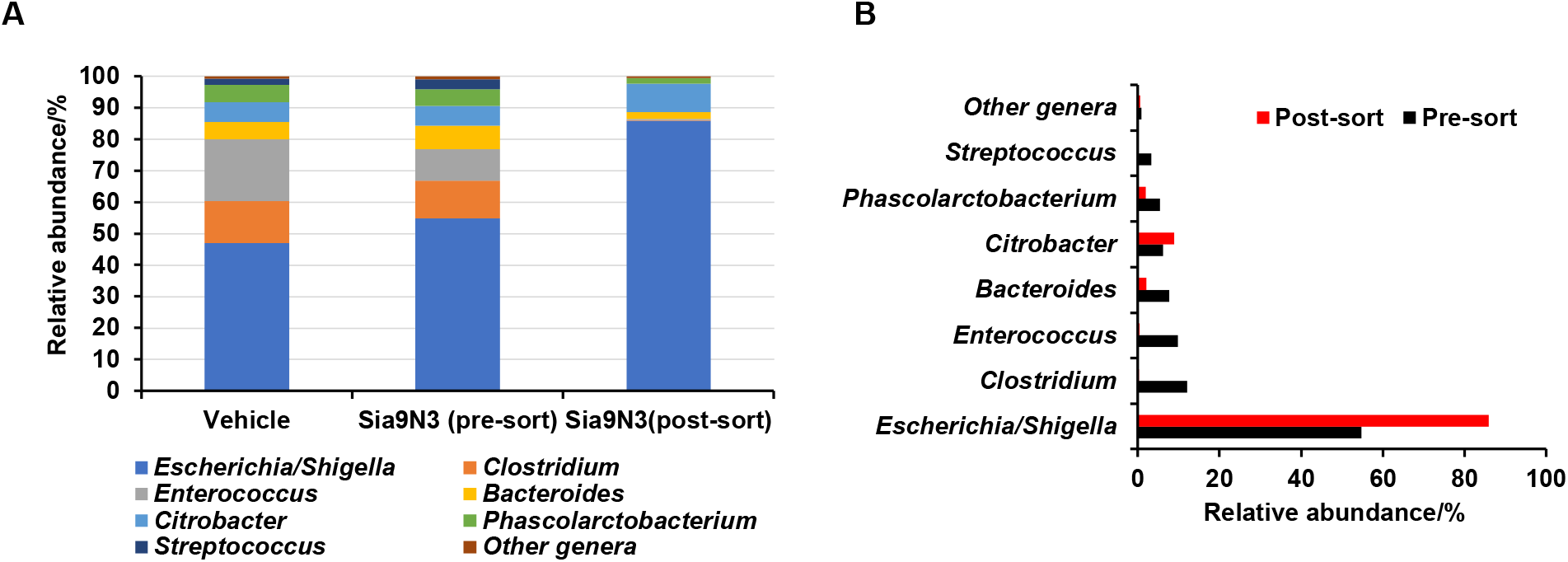
16S rDNA metagenomic sequencing analyses. **A.** H2 microbiome sample was cultured with or without 200 μM Sia9N_3_, which does not significantly alter taxonomic composition of 20-h H2 microbiome culture. Sia9N_3_ presenting bacteria from H2 microbiome culture were labeled with fluorescence and isolated with FACS. **A and B.** 16S rDNA sequencing showed that the *Escherichia/Shigella genus* was significantly enriched after cell sorting.

To identify the bacterial genome of the Sia9N_3_-presenting H2 *Escherichia*, we randomly selected 10 colonies from an H2 culture plate followed by Sia9N_3_ metabolic labeling assay. Out of 10 colonies screened, liquid culture in Gifu media of colony 2 passage 1 yielded nearly 100% incorporation of Sia9N_3_ (C2-P1, **Fig. 5A**). Furthermore, 3 randomly picked colonies from a C2 culture plate (C2 passage-2, C2-P2-1, −2, and −3) also showed 100% Sia9N_3_ incorporation, suggesting that the Sia9N_3_ catabolic genes were readily passaged to, and active in, the daughter generations (**Fig. 5B**). The ZH-C2 colony was subjected to shotgun whole-genome sequencing in an attempt to identify the *Escherichia* bacterial strain.

**Fig. 5.**
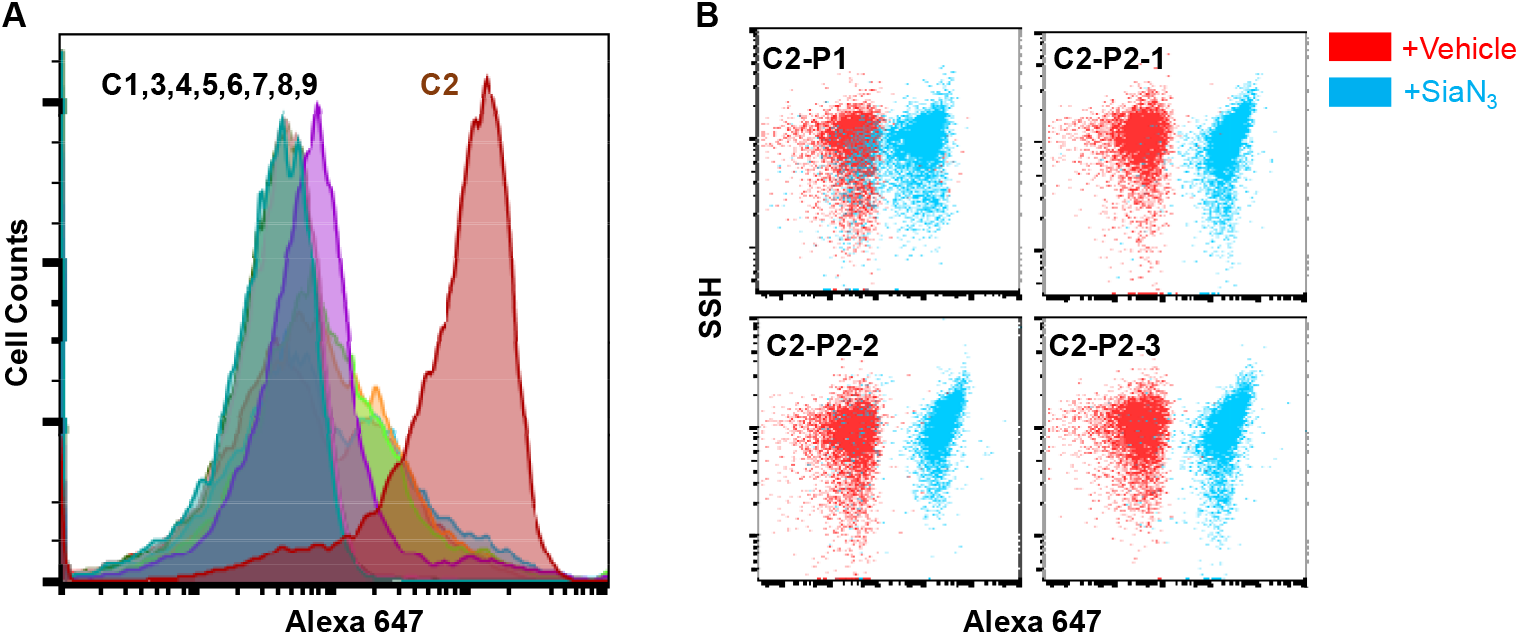
Isolation and passaging of the Sia9N_3_ incorporating bacterial strain from H2 microbiota. **A.** Colony screening of H2 microbiota isolated one Sia9N_3_ incorporating bacterial strain. **B.** The Sia9N_3_ incorporating feature is readily passaged to daughter generations.

The genome of our newly isolated bacteria (ZH-C2) cannot be fully mapped to any previously sequenced *Escherichia* bacteria in the NCBI RefSeq database, but is most similar to pathogenic *E. coli* strains UMEA 3174-1 (accession PRJNA186306) and VR50 (accession PRJEA61445) with average nucleotide identities of 99.85% and 99.76%, respectively (**Fig. 6A** and **Supplemental Table, tab 1**). Our *E. coli* ZH-C2 strain has a 5.23 kb genome that is similar in size to UMEA 3174-1 and VR50 and is much larger than the 4.64 kb genome of the prototypical nonpathogenic strain K-12 MG1655 (MG1655, accession PRJNA647163) (**Supplemental Table, tab 1**). ZH-C2 shares over 90% identity in sequence to most MG1655 genes; however, ZH-C2 consists of 1,111 missing or low-homology genes compared to MG1655, while MG1655 has 347 missing or low-homology genes relative to ZH-C2 (**Fig. 6A** and **Supplemental Table, tab 2**). The majority of gene products have no identified function; however, GO term analyses demonstrate ZH-C2 encodes for an increased diversity and number of additional proteins relative to MG1655 involved in DNA repair and replication, polysaccharide transport, viron assembly, as well as type II secretion (**Fig. 6B** and **Supplemental Table, tab 3**). With respect to molecular function, a diverse set of additional genes are encoded in ZH-C2, notably predicted cobalamin binding, ATPase, transferase, and porin activities (**Fig. 6C** and **Supplemental Table, tab 3**) with a significantly larger collection of ZH-C2 proteins likely localized to the outer membrane compared to MG1655 (**Fig. 6D** and **Supplemental Table, tab 3**). Conversely, the MG1655 genome consists of a number of additional genes associated with DNA-mediated recombination and integration, as well as cell adhesion, DNA binding and DNA recombinase activities (**Fig. 6B-D** and **Supplemental Table, tab 3**).

**Fig. 6.**
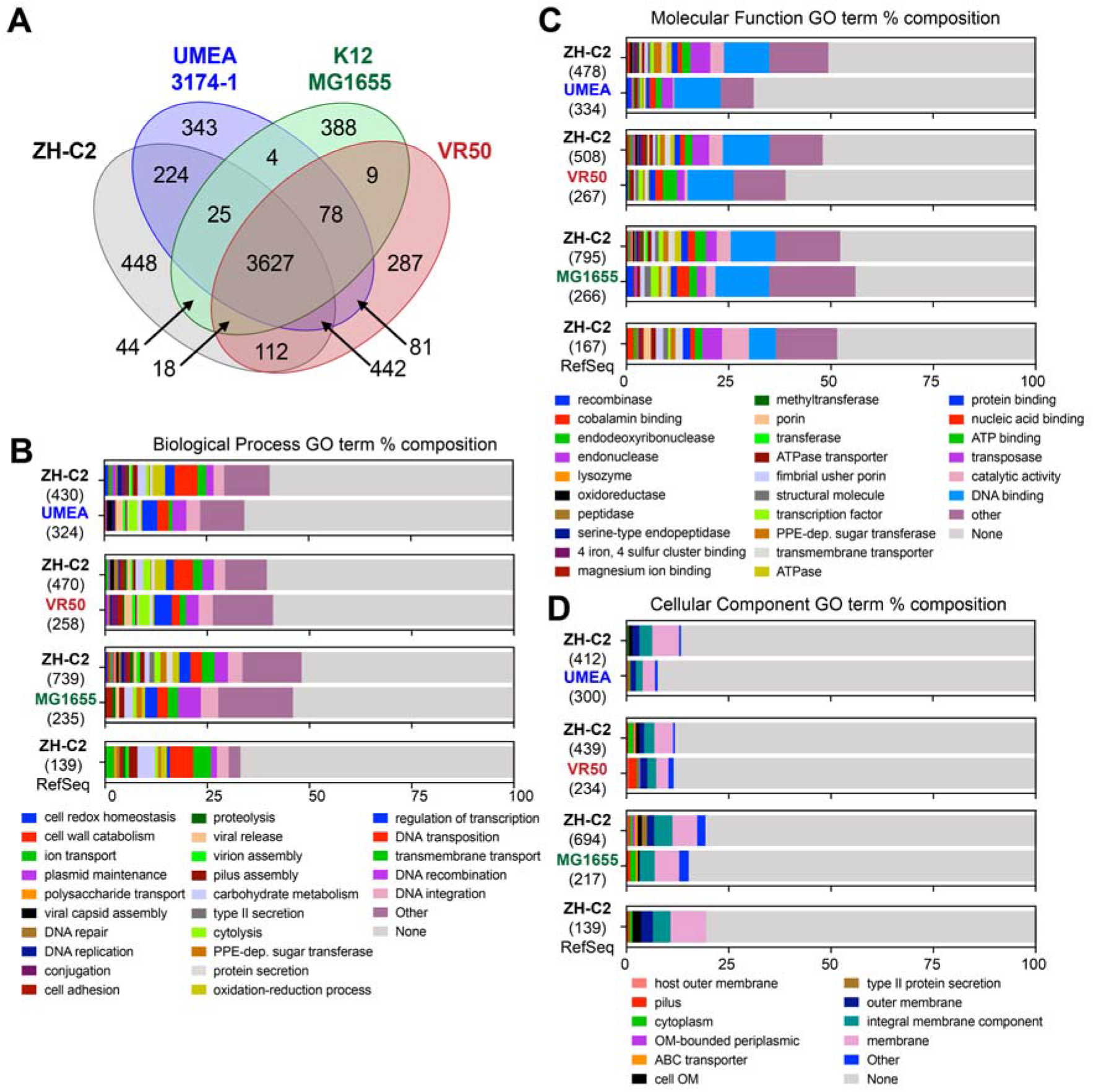
Comparison of encoded *E. coli* ZH-C2 genes to reference non-pathogenic and pathogenic *E. coli*. Venn diagram of CD-HIT analysis depicting conservation and differences of gene products between *E. coli* strains ZH-C2, K-12 MG1655, UMEA 3174-1, and VR50 (**A**). Protein sequences were clustered at CD-HIT sequence similarity cut-off of 90%. Biological process (**B**), molecular function (**C**), and cellular component (**D**) GO Terms of unique genes (less than 70% coverage and/pr less than 90% identity) found in ZH-2 compared to MG1655, UMEA 3174-1, VR50, and the RefSeq reference genome collection. The number of genes used for each GO term analysis is in parentheses. GO Terms listed as “other” include <2 genes and are expanded in the **Supplementary Table.**

MG1655 encodes for the sialic acid catabolizing genes, including Nan A, C, E, K, M, the regulatory gene NanR, and sialic acid transporter gene NanT. However, as for *E. coli* strains UMEA 3174-1 and VR50, ZH-C2 encodes for: 1) NeuA, the cytidine monophosphate (CMP)-sialic acid synthetase which produces the sialyltransferase donor CMP-sialic acid; and 2) NeuS, the polysialyltransferase which synthesizes capsular polysialic acids from CMP-sialic acid (29). The lack of the NeuA and NeuS genes highlights the inability of MG1655 to activate and present Sia9N_3_ on its cell surface (**Fig. S4**). Moreover, the sugar-phosphate isomerase KpsF/GutQ (accession WP_001296394, ZH-C2_01729), known to be associated with polysialic acid deposition onto the capsule is also identified in the ZH-C2 genome (with 100% identity to UMEA 3174-1 and VR50) but not in MG1655 (37). Altogether, the genomic analyses suggest that the cell-surface fluorescence labeling we observed is caused by deposition of polySia9N_3_ on the bacterial capsule.

Despite being isolated from a healthy human fecal sample, our ZH-C2 strain may be a pathogenic/pathobiont *E. coli*. The two most similar genomes, UMEA 3174-1 and VR50, were both isolated from urinary tract infections (**Supplemental Table, tab 1**). Established pathogenic genes, such as the type III secretion system are conserved across ZH-C2, UMEA 3174-1, and VR50 (38). Notwithstanding the high conservation, notable differences were also observed. ZH-C2 has 628 and 680 low homology/unique genes relative to UMEA 3174-1 and VR50, respectively. Conversely, UMEA and VR50 have 507 and 393 low homology/unique genes, respectively, to ZH-C2 as determined by sequences with <90% identity and/or covering <70% of the sequence length (**Supplemental Table, tabs 4,5**). Interestingly, there is an enrichment of predicted GO terms associated with DNA-mediated transposition, oxidation-reduction processes, pilus assembly, and carbohydrate metabolic activities in ZH-C2 relative to both UMEA 3174-1 and VR50, as well as a localization of proteins to the membrane (**Fig. 6B-D** and **Supplemental Table, tab 3**). Those missing functions in ZH-C2 found in UMEA 3174-1 and VR50, include transcriptional regulation, viral release, nucleic acid binding, and recombinase activity (**Fig. 6B-D** and **Supplemental Table, tab 3**).

With respect to all available non-redundant RefSeq reference genomes, ZH-C2 has 207 low homology or unique proteins with approximately 65 sequences consisting of <100 amino acids (**Fig. 6B-D** and **Supplemental Table, tab 6**). Almost all of these sequences have been identified in one or more of the other >20K *E. coli* strains that have been sequenced to date and are in the NCBI RefSeq/GenBank database. However, ZH-C2 encodes for ZH-C2_04397 and ZH-C2_04599 that are 1,174 and 475 amino acids long, respectively, and have not been observed previously in any *E. coli* strain. ZH-C2_04397 is a multi-subunit protein consisting of an N-terminal autoinducer-2 kinase, an autotransporter barrel domain-containing lipoprotein, and a C-terminal HipA-like Ser/Thr kinase (**Fig. S5**) (39,40). While most *E. coli* encode for a linked autotransporter lipoprotein and C-terminal HipA-like Ser/Thr kinase, only one other *E. coli* has an N-terminal autoinducer-2 kinase connected to the autotransporter lipoprotein (LsrK in *E. coli* KTE98, accession EOV99460.1). However, this *E. coli* KTE98 gene lacks the C-terminal HipA-like Ser/Thr kinase. Most *E. coli* do encode for an autoinducer-2 kinase as a single gene. ZH-C2_04599 also is a multidomain gene consisting of an N-terminal transposase and a C-terminal EamA transporter (**Fig. S6**) (41). Again, both domains are well-represented across *E. coli* as single genes. The biological importance of these multi-functional connected domains will be subject to future investigations.

## Discussion

Azide-modified glucosamine, mannosamine, fucose, and 3-deoxy-D-manno-oct-2-ulosonic acid (Kdo) have been used to metabolically label bacterial cells *in vitro* and *in vivo* (19,20,42). Here, we demonstrate the application of azide-modified N-acetylneuraminic acid to selectively label and track Neu5Ac-presenting bacteria from a complex human fecal microbiome sample for the first time. Using flow cytometry and metagenomic sequencing, we identified a new strain of *E. coli* that can present environmental Neu5Ac on its surface via the NanT-NeuA-NeuS pathway. Our results also suggest a dynamic process by which select distal gut bacteria can exploit environmental sialic acid for self-decoration that, in turn, is likely subject to removal by sialidase activity provided by other microbiome constituents (*e.g., B. thetaiotaomicron*).

The Sia9N_3_-presenting activity of our new *E. coli* ZH-C2 strain is only detected in sample H2. Additional studies will investigate if this new *E. coli* is missing from the H1 and H3 cultures or present with a silenced NanT-NeuA-NeuS pathway during the *ex vivo* culture. Previous studies suggest both possibilities, as the gut microbiome is regulated by diet and crosstalk amongst bacteria as well as the host and suggests a dynamic composition of different bacterial strains over time (43,44). Conversely, certain bacteria, including *Bacteroides fragilis*, are able to switch capsular polysaccharide structures enabling the colonization specific intestinal niches (45). The gradual decrease in *E. coli* Sia9N_3_ surface expression over time will also be further interrogated. We anticipate that the loss of Sia9N_3_ is caused by other commensal organisms that gain prominence over time in the *ex vivo* culture and remove the *E. coli* poly-SA as a biologically relevant mechanism for host recognition and clearance. Additionally, surface-expressed poly-SA may be harvested by the parental *E. coli* for growth, as environmental conditions and nutrient availability shift over time.

Distal gut bacteria can degrade and ferment complex carbohydrates to provide an energy source and/or immunomodulatory molecules for themselves, the host, and/or other gut bacteria (1,32,46,47). In return, host cells can produce glycans which can provide nutritional advantages for certain microbes (48). Such mutualistic relationships make the tracking and study of carbohydrate catabolism by individual bacteria in complex microbiomes complicated and challenging. Consistent with previous studies, we found that cultured microbiome samples do not fully represent the taxonomic diversity of the original source due to growth advantages of certain bacteria under anaerobic systems with limited nutritional sources from the selected media (31,49). Notwithstanding, our approach enriched for a new SA-presenting strain of *Escherichia coli* isolated from a human gut microbiome. We posit that SA-presenting bacteria can be identified from different microbial sources, including respiratory, urinary, and vaginal microbiomes in both health and disease (*i.e.,* detection of pathogens in blood and stool clinical samples). Additionally, the scavenging of environmental sialic acid is associated with pathogenic bacterial virulence (50) and proteins involved in the process may represent new targets for the discovery of antibacterial small molecules.

## Conclusions

Our integrated assay system detects sialic acid-presenting bacteria in real-time from complex microbiome samples. We envision that this labeling and tracking strategy can assist in identifying new SA-presenting bacteria and developing diagnostic tools and drug discovery models. Furthermore, this workflow may be used to investigate the installation of other environmental glycans by microbiota.

## Supplemental Information

Detailed methods and reagents used in this study are included in the supplemental information.

## Acknowledgements

We thank Jenny Cornell, Jessica Ledesma, and Steven Robert Head at Next Generation Sequencing Core at The Scripps Research Institute for 16S rDNA and whole-genome sequencing experiments. We thank Padmaja Natarajan and Alain Domissy at Center for Computational Biology at The Scripps Research Institute for analyzing the 16S rDNA sequencing data. We thank Qiyun Zhu at the University of California, San Diego for assistance with the whole-genome sequencing data analysis. This work was supported by NIH award R21 AI139744 to D.W.W. and Scripps Research Translational Institute, an NIH-NCATS Clinical and Translational Science Award (CTSA; UL1 TR002550) to W.P. and A.T.

## Author contributions

Z.H. and D.W.W. designed the research; Z.H., P.T.B., V.F.V., and J.R.T. performed the experiments; Z.H., W.P., P.T.B., A.T., and D.W.W analyzed the data; Z.H. and D.W.W. composed the paper and all authors edited and approved its contents.

## Declaration of interests

The authors declare no conflict of interest.

## Notes

### Competing Interest Statement

The authors have declared no competing interest.

