## Supplemental Information for "Identification of an N-acetylneuraminic acid-presenting bacteria isolated from a healthy human microbiome"

- **EXPERIMENTAL MODEL AND SUBJECT DETAILS**

**Preparation of Gifu culture media and agar plate**

Gifu Anaerobic Broth, Modified (GAM, M2079), was purchased from HIMEDIA. 8.4 g of powder was dissolved in 1 L milli-Q water, followed by autoclaving at 115 °C for 15 min. Sterile media was cooled down and filtered through a 0.22 μm filter before use. Gifu-agar plates contain same concentration of Gifu media plus 1.2% agar.

**Fecal Microbiome sample collection and culture**

Three human fecal samples (H1, H2, and H3) were collected from one healthy donor longitudinally at day 1, 18, and 80. Immediately after collection, fecal samples were transported within an anaerobic bag on dry ice into an anaerobic chamber (Coy Laboratory Products) filled with 3% hydrogen and 97% $N_2$ within 10 min. Fecal samples were then re-suspended in filtered phosphate buffer saline (PBS) buffer, pH 7.4. Large non-bacterial particles were settled down and the cloudy supernatant was carefully aspirated out. Bacterial cells were then centrifuged down at 16000xg for 5 min and washed with PBS 3 times. Fecal bacteria were cultured in the Gifu media in anaerobic atmosphere at 37 °C with an initial optical density at 600 nm ($OD_{600}$) of 0.005. The rest of bacterial cells were re-suspended in 20% glycerol/PBS solution at an $OD_{600}$ of 10, flash frozen, and stored at -80 °C for future use.

- **METHOD DETAILS**

**Synthesis, purification, and characterization of Neu5Ac9N$_3$**

ACS reagent grade organic solvents were purchased from Sigma Aldrich and were used directly if not specified. N-Acetyl-9-azido-9-deoxy-neuraminic acid (Neu5Ac9N$_3$/Sia9N$_3$) was synthesized, purified, and characterized as previously reported (1). Briefly, Neu5Ac (3 g, 9.6 mmol) was mixed with Dowex 50Wx8-200-400 (H) (2g, Alfa Aesar, L13922) in 50 mL methanol, followed by overnight stirring at room temperature. The resin was removed by filtration over Celite and the filtrate was concentrated under reduced pressure to 15 mL. 15 mL of ice-cold diethyl ether was layered above the concentrated methanolic solution, and the mixture was chilled at 4 °C overnight to yield a pale yellow crystalline solid. Supernatant was carefully decanted, and the pellet was carefully rinsed with ice cold diethyl ether twice, then air dried to give 2.2 g Neu5AcOMe as a white crystalline powder. Neu5AcOMe (2.2 g, 6.8 mmol) was dissolved in 20 mL of anhydrous pyridine, cooled in an ice bath, then treated with *p*-toluenesulfonyl chloride (2.35 g, 12.3 mmol; freshly purified by recrystallization from chloroform and petroleum ether). The mixture was stirred over an ice bath for 30 min, followed by overnight stirring at room temperature. Pyridine was removed by rotary evaporation to form syrup-like crude, which was purified with flash chromatography (EtoAc:MeOH=20:1~20:5), dried, to yield 0.6 g Tosyl-Neu5AcOMe as white solid. Tosyl-Neu5AcOMe (0.6 g, 1.26 mmol) and $NaN_3$ (0.32 g) were suspended in 10 mL of acetone:water (3:1) solution followed by overnight stirring at 70 °C. The crude was dried, acidified with Dowex 50Wx8-200-400 (H), and purified with flash chromatography (60:40~25:75 DCM:MeOH), dried to yield 0.31 g pure final product Neu5Ac9N$_3$/Sia9N$_3$.

**Sequential fluorescence labeling of bacterial cells and flow cytometry**

Fecal microbes cultured with or without Sia9N$_3$ were collected by centrifugation at 16000xg and washed with filtered PBS for 3 times. Bacterial cells were then re-suspended to an $OD_{600}$ of 1.0, treated with

DBCO-Biotin for 2 h at room temperature in anaerobic atmosphere. The bacterial cells were washed with PBS for 3 times with centrifugation at 16000xg for 5 min, followed by streptavidin Alexa 647 staining at 1:200 dilution for 1 h with $OD_{600}$ at 1.0. Stained bacterial cells were washed with PBS for 3 times, followed by analyzing on a NovoCyte 3000 with NovoSampler Pro model. Singlet cells were selected by plotting SSC-H and FSC-H versus SSC-A and FSC-A, respectively, followed by gated on Alexa 647 channel. 15000 total events were collected for data analysis with the percentage of singlets at ~90%.

**Fluorescence microscopy**

Fluorescently labeled bacterial cells were spotted on a glass slip, airdried for 15 min, and covered with a cover slip with ~1 nm thickness. Confocal microscopy was performed on a Zeiss LSM 880 Airyscan confocal laser scanning microscope. Imaging analyses were processed with FlowJo_v10.

**Expression and purification of sialidase BT0455**

The gene fragment encodes for the *Bacteroides thetaiotaomicron* sialidase BT0455 (Uniprot ID: Q8AAK9) was ordered from Integrated DNA Technologies (IDT, with the N-terminal signal peptide removed. The BT0455 encoding sequence was cloned into a customized pT7HMT vector (2), followed by expression in BL21(DE3) cells as an N-terminal $His_6$ tag fusion. BT0455 was purified from bacterial cell lysate with a nickel-nitrilotriacetic acid (Ni-NTA) affinity column, followed by size-exclusion chromatography. Finally, BT0455 was aliquoted and stored at -80 $^{o}$C in a storage buffer containing 20 mM Tris-HCl and 100 mM NaCl with pH at 8.0.

**Sialidase activity assay**

14-66 h after fecal microbiota culture, the bacterial cells were pelleted and the supernatants (cultured media) were assayed for sialidase activity. Specifically, a 200 uL PBS mixture containing 500 pM purified BT0455 sialidase or cultured media and 100 $\mu$M turn-on fluorescent substrate 4-methylumbelliferyl-$\alpha$-D-N-acetylneuraminic acid (4-MUNANA) was pre-incubate for 60 s, followed by fluorescence measurement on a microplate reader (excitation 365 nm, emission 450 nm). The fluorescence intensities were measured every 60 s over the linear range of enzymatic reaction. The presented data is a representative of three individual activity assays to ensure reproducibility.

**FACS**

Fluorescence activated cell sorting (FACS) of fluorescently labeled bacteria from H2 microbiota culture was performed on a Sony MA900 multi-application cell sorter. Singlets cells were selected by plotting SSC-H and FSC-H versus SSC-A and FSC-A, respectively. Approximately $1x10^7$ fluorescently labeled bacterial cells were sorted and collected for 16S rDNA sequencing analyses.

**DNA extraction and sequencing analysis**

Pre-sorted or post-sorted bacteria were subjected to DNA extraction with the ENZA bacterial DNA extraction kit (Omega biotek, D3350) based on manufacture's protocol. 5-22 ng DNA was used with the NEXTFLEX 16S V4 Amplicon-Seq Kit 2.0 library prep kit following manufacturer's recommended protocol with between 11 - 20 PCR cycles depending on sample input. PCR amplicons were all pooled and gel purified to select the target amplicon region. Libraries were loaded onto a flowcell and sequenced on an Illumina MiSeq sequencer to generate 2 x 300-bp paired-end reads. Generated reads were then analyzed with Illumina Basespace online software for microbiome taxonomic annotation.

**Colony screening**

Human microbiota H2 was plated on a Gifu-agar plate for overnight culture in an anaerobic atmosphere. 10 colonies were randomly picked and inoculated in 1 mL Gifu-Sia9$N_3$ media for 20 h anaerobic culture at 37 °C. Then, bacteria were tested for fluorescence labeling with flow cytometry as described. Bacteria cultured from C2 colony (C2 passage-1, C2-P1) showed nearly 100% fluorescence labeling. To test the homogeneity of C2 and C2-P1 bacteria, C2-P1 bacteria were cultured on a Gifu-agar plate and 3 colonies were randomly picked and inoculated in Gifu-Sia9$N_3$ media followed by cultivation and fluorescence labeling experiments. As a result, 100% bacteria cultured from all three colonies (C2-P2-1~3) were fluorescently labeled.

**Shotgun whole-genome sequencing**

Colony C2-P2-2 and C2-P2-3 were cultured in 6 mL Gifu media, respectively. Bacterial cells were then pelleted and washed, and bacterial DNA was extracted with ENZA bacterial DNA extraction kit (Omega biotek, D3350) based on manufacture's protocol. For genomic preps, DNA was sheared to ~500 bp size using a Covaris S2 Focused-ultrasonicator, after which 10 ng of sheared DNA was used to make sequencing libraries with the NEBNext® Ultra™ II DNA Library Prep Kit following manufacturer's recommended protocol with 9 cycles of PCR. Libraries were size selected using AmpureXP beads to 500-700 bp, loaded onto a flowcell, and sequenced on an Illumina MiSeq sequencer to generate 2 x 300-bp paired-end reads.

**Whole-genome assembly of ZH-C2**

The reads for ZH-C2 were processed by Trimmomatic 0.36 (3) to remove the Illumina adapters and parts of the reads with low quality using the following filters: ILLUMINACLIP:<path-to-adapters>/TruSeq3-PE-2.fa:2:30:10 LEADING:30 SLIDINGWINDOW:10:20 MINLEN:100. The trimmed reads were then assembled using SPAdes 3.13.0 (4) with the following k values: 21, 33, 55, 77, 99, 127. Next, the blastn command of BLAST 2.7.1+ (5) with -max_target seqs 1 was used to align the assembled ZH-C2 contigs against the NCBI RefSeq genome database (https://www.ncbi.nlm.nih.gov/genome). Two custom Perl scripts binned the voluminous BLAST output by species and strain. The average nucleotide identity (ANI) between ZH-C2 and the top-scoring strains was computed using blastn with -best_hit_overhang 0.1 -best_hit_score_edge 0.1 and another custom Perl script to determine the closest *E. coli* genomic strains: UMEA 3174-1 (closest overall), VR50 (second closest), and K-12 MG1655 (closest of three reference genomes). Some 11% of the ZH-C2 contigs did not align with *E. coli.* These were presumed to be from contaminants and were removed from the assembled genome along with another 5% of the contigs that were shorter than 200 bp to comply with guidelines for submission to GenBank.

**Identification of genes and their protein products**

The proteins produced by UMEA 3174-1, VR50, and K-12 MG1655 are available from the RefSeq database, For ZH-C2, Prokka 1.14.6 (6) was used to identify the genes and associated proteins. Many were identified only as "hypothetical proteins". To get further information on these as well as more consistent naming, a database of RefSeq bacterial proteins was generated using the commands given at https://dmnfarrell.github.io/bioinformatics/local-refseq-db. The blastp command of BLAST with -max_target_seqs 1 was then used to align the proteins from Prokka against the new database. This provided additional information on more than half of the hypothetical proteins examined; 19 proteins from Prokka were not found in RefSeq.

**Identification and removal of duplicate proteins**

To check for the presence of duplicate proteins in a single genome, the proteins in each genome were aligned against themselves using blastp with -max_target_seqs 15. Most matches had less than 100% identity and 100% coverage, and these were ignored. The remaining perfect matches were mostly unique but also revealed 6 duplicate proteins in ZH-C2 from Prokka and 41 duplicate proteins in K-12 MG1655 from RefSeq. These duplicates were removed for subsequent analysis. No duplicates were found in the proteins for UMEA 3174-1 or VR50 from RefSeq.

**Comparison of proteins**

The proteins in ZH-C2 were compared with those in each of the three other genomes of interest, and vice versa, using blastp. Even though -max_target_seqs 1 was specified, some proteins had more than one match with varying lengths. Thus, a second pass was made through the blastp output retaining only the longest match when there was more than one. Finally, the matches for each of the six pairwise comparisons were sorted into four groups corresponding to (1) perfect matches between the two genomes, (2) high-similarity matches with at least 90% amino acid identity covering at least 70% of the protein length, (3) low-similarity matches that did not meet the previous criteria, and (4) non-matches.

Venn diagrams were generated using output from CD-HIT (4.8.1) clustering of 4 combined proteomes (ZH-C2, MG1655, UMEA 3174-1, VR50) (7,8). The following command line input was used to generate desired cluster files: "cd-hit -i inputfile.fasta -o outputfile -c %cutoff -g 1 -d 0" where "%cutoff" is the percent similarity cut-off used to define a cluster. In our analysis, we typically set percent similarity cut-offs at 65%, 75%, 85%, 95%, and 99%. CD-HIT output was organized into tables using the Python Pandas package and Venn diagrams were generated using venn/pyvenn packages.

Gene Ontology (GO) terms were generated using InterProScan (5.40-77.0) (9). Briefly, protein sequences of interest were concatenated into a protein fasta file then submitted for InterProScan analysis using the following command line input: "./interproscsan.sh -i input_file.fasta -f tsv -dp -goterms". Output tab separated files were organized using the Python Pandas package and bar plots were generated in Matplotlib. GO terms were mapped to their name spaces using the comprehensive "go.obo" listing available at geneontology.org (10). GO terms were organized such that all protein sequences contributed at least 1 count to each of the 3 "Biological Process," "Molecular Function," and "Cellular Component" GO namespaces. If sequences could not be annotated, 1 count of "None" was contributed to each of the name spaces. For sequences annotated multiple times, each annotation contributed 1 count to the GO term's respective namespace.

- **QUANTIFICATION AND STATISTICAL ANALYSIS**

In **Figure 3C**, the mean RFU$\pm$standard deviation of a triplicate experiment is shown.

**Supplementary Figures and Tables**

**Fig. S1.** Dose-dependent labeling of H2 cultured microbiome sample.

**Fig. S2.** Sia9$N_3$ metabolic labeling of microbiota H2 competed with Neu5Ac and removed by a microbial sialidase BT-0455.

**Fig. S3.** 16S rDNA sequencing of primary and cultured microbiome samples.

**Fig. S4.** Sia9$N_3$ incorporation by *E. coli*-K12 (MG1655) and the newly isolated *E. coli* strain.

**Fig. S5.** Sequence alignment of ZH-C2_04397 to other *E. coli* proteins.

**Fig. S6.** Sequence alignment of ZH-C2_04599 to other *E. coli* proteins.

**Supp. Table.** 6 tabs

**Supplemental Table legend: Additional bioinformatic data analyses of *E. coli* genome composition.** Related to Figure 6. Tab 1: *E. coli* genome assembly statistics and comparisons to reference strains. Tab 2: List of low-homology and unique *E. coli* ZH-C2 genes relative to K-12 MG1655. Tab 3: Biological process, molecular function, and cellular component GO Terms associated with unique *E. coli* genomes. Tab 4: List of low-homology and unique *E. coli* ZH-C2 genes relative to UMEA 3174-1. Tab 5: List of low-homology and unique *E. coli* ZH-C2 genes relative to VR50. Tab 6: List of low-homology and unique *E. coli* ZH-C2 genes relative RefSeq collection. Low homology hit proteins are best match spans with <70% of query protein length and/or <90% positive identity rate.

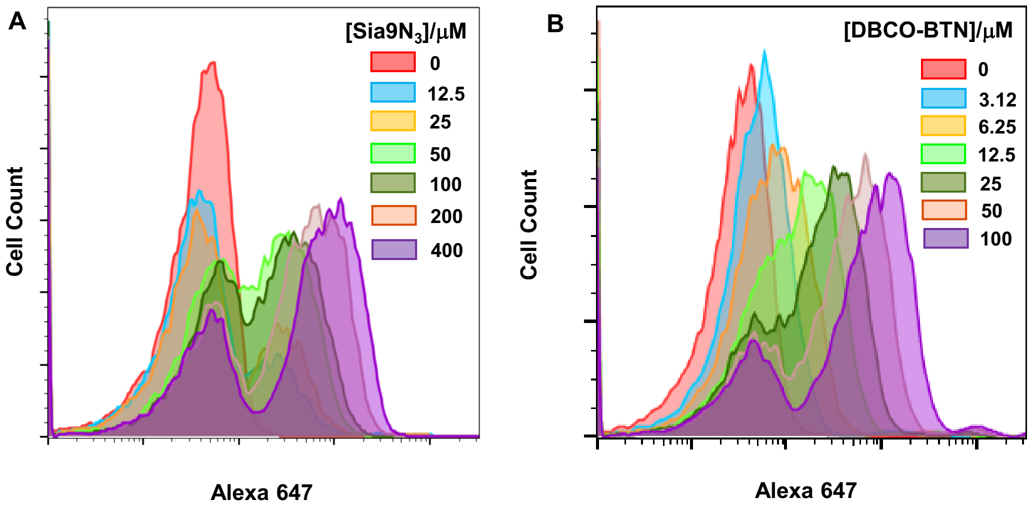


**Figure S1.** Sia9$N_3$ (**A**) and DBCO-BTN (**B**) dose-dependent labeling of H2 cultured microbiome sample.

**A.**

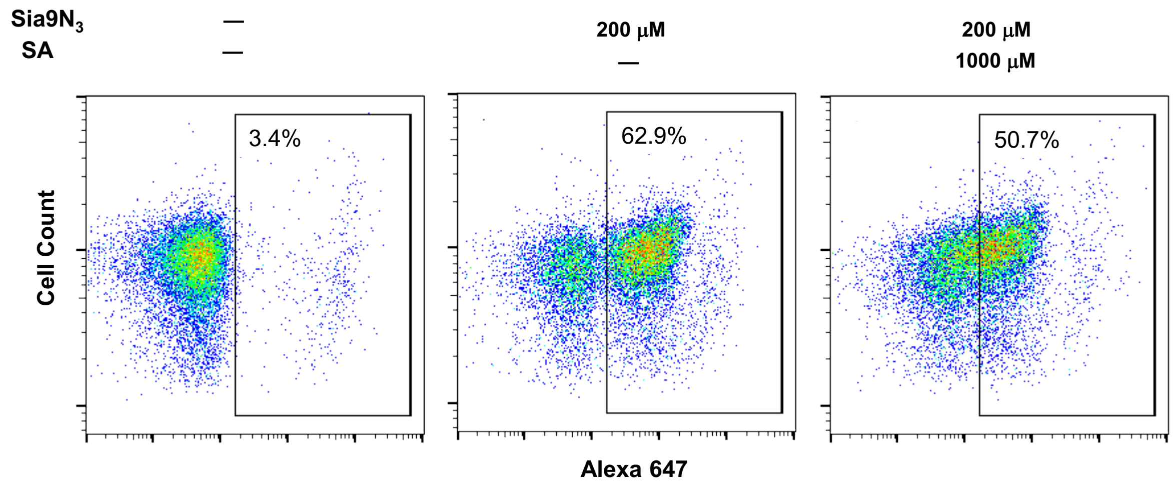


**B.**

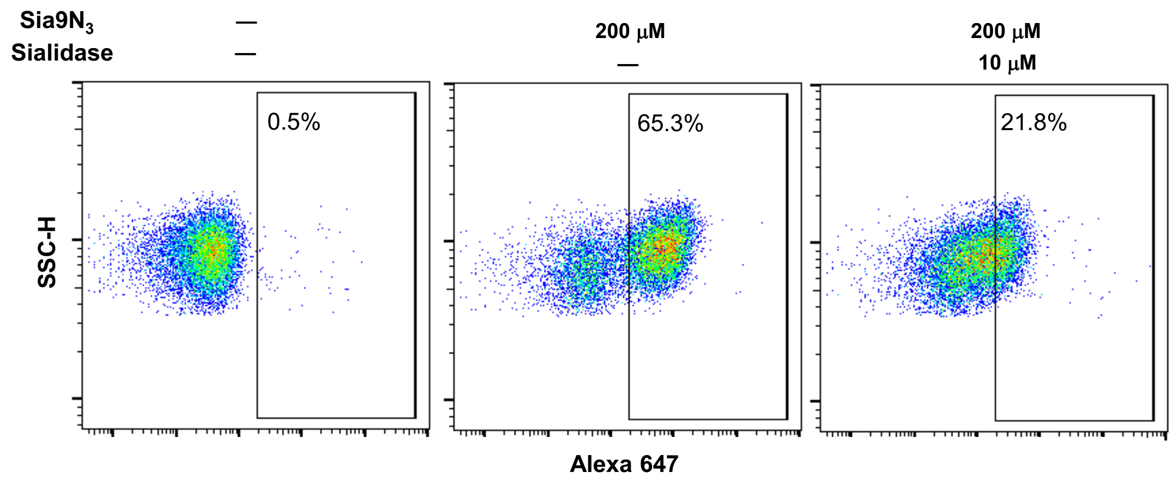


**Figure S2. A.** Sia9$N_3$ metabolic labeling of microbiome H2 competed with Neu5Ac. **B.** Sia9$N_3$ metabolic labeling is partially removed by a microbial sialidase BT-0455.

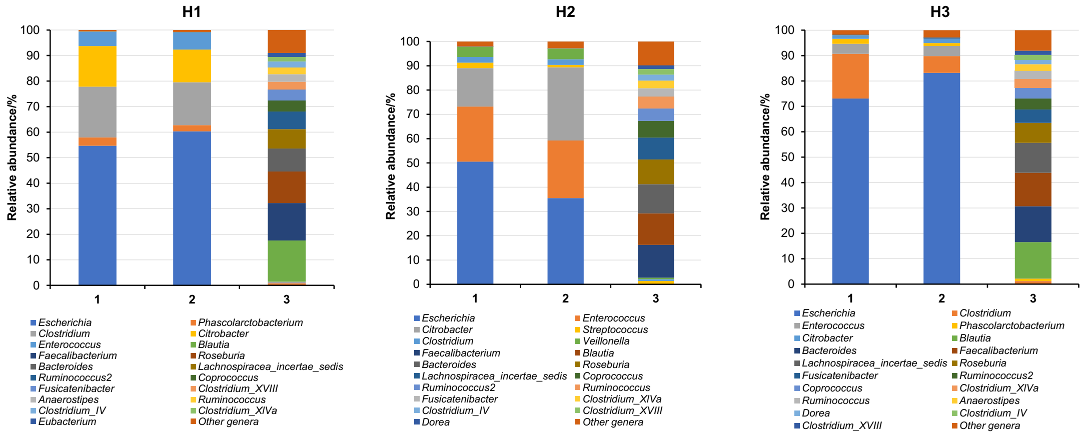


**Figure S3. 16S rDNA sequencing revealed taxonomic composition of cultured and primary microbiomes in the presence and absence of 200 μM Sia9N$_3$.** Cultured human microbiome samples H1-H3 were all tested, column 1: microbiome cultured without Sia9N$_3$, column 2: microbiome cultured with Sia9N$_3$, 3: uncultured fecal microbiome samples.

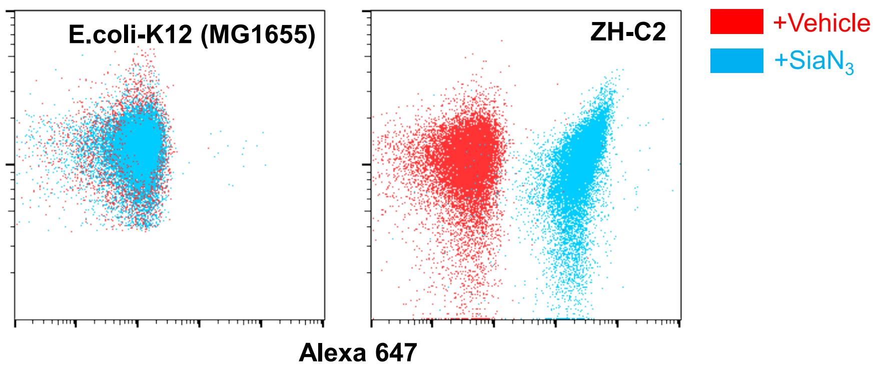


**Figure S4.** Test of Sia9$N_3$ incorporation by *E. coli*-K12 (MG1655) and the newly isolated *E. coli* strain ZH-C2.

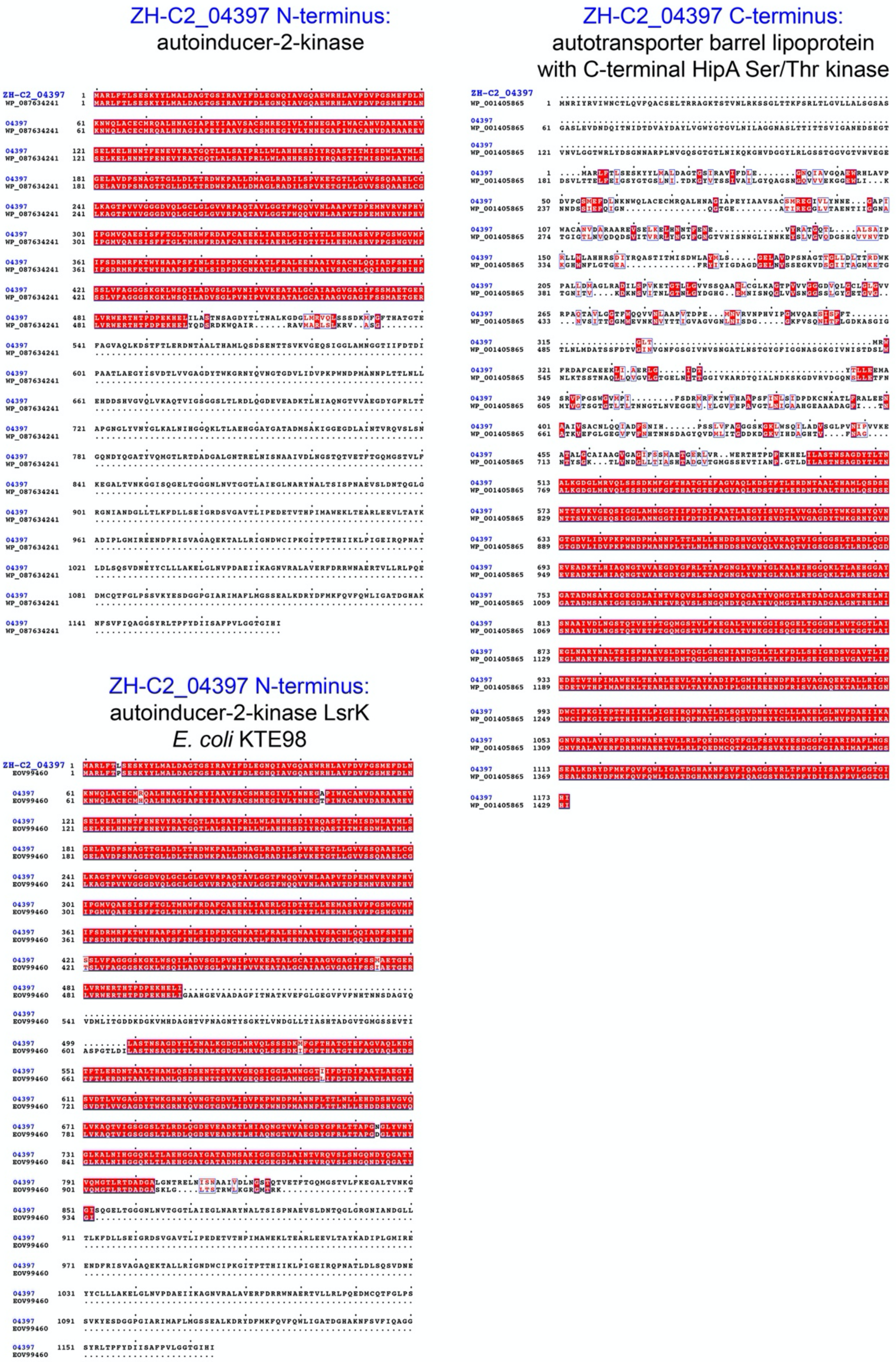


**Figure S5.** Unique multi-domain ZH-C2_04397 with an N-terminal transposase, autotransporter barrel lipoprotein, and C-terminal HipA-like Ser/Thr kinase aligns to single-domain *E. coli* proteins.

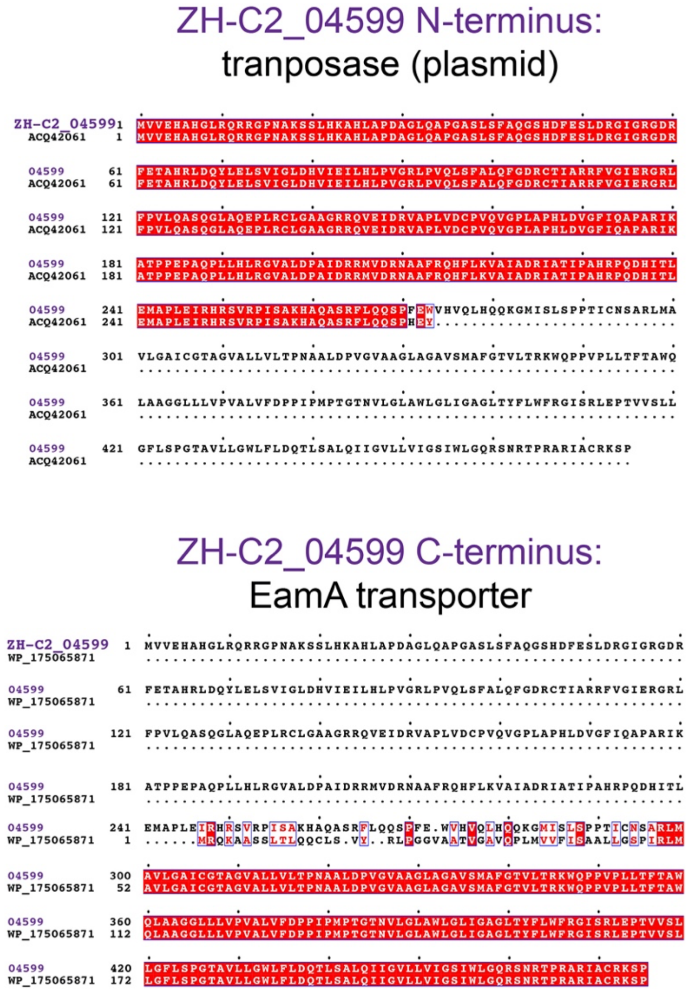


**Figure S6.** Unique multi-domain ZH-C2_04599 with an N-terminal transposase and C-terminal EamA transporter aligns to single-domain *E. coli* proteins.
